# A machine-learning approach to human footprint index estimation with applications to sustainable development

**DOI:** 10.1101/2020.09.06.284414

**Authors:** Patrick W. Keys, Elizabeth A. Barnes, Neil H. Carter

## Abstract

The human footprint index is an extensively used tool for interpreting the accelerating pressure of humanity on Earth. Up to now, the process of creating the human footprint index has required significant data and modeling, and updated versions of the index often lag the present day by many years. Here we introduce a near-present, global-scale machine learning-based human footprint index (ml-HFI) which is capable of routine update using satellite imagery alone. We present the most up-to-date map of the human footprint index, and document changes in human pressure during the past 20 years (2000 to 2019). Moreover, we demonstrate its utility as a monitoring tool for the United Nations Sustainable Development Goal 15 (SDG15), “Life on Land”, which aims to foster sustainable development while conserving biodiversity. We identify 43 countries that are making progress toward SDG15 while also experiencing increases in their ml-HFI. We examine a subset of these in the context of conservation policies that may or may not enable continued progress toward SDG15. This has immediate policy relevance, since the majority of countries globally are not on track to achieve Goal 15 by the declared deadline of 2030. Moving forward, the ml-HFI may be used for ongoing monitoring and evaluation support toward the twin goals of fostering a thriving society and global Earth system.

## Introduction

The human footprint index (HFI) represents one of the most important tools for interpreting human pressure on the landscape (Venter *et al* 2016b). A dimensionless metric which captures the extent of human influence on the terrestrial surface (Fig. 1a), the HFI is distinct from many land-use metrics in that it captures the total influence of human existence on a given location in a single index, rather than categorizing individual land use types (Riggio *et al* 2020). The applications to which the HFI is put to use are manifold (Di Marco *et al* 2018, Watson *et al* 2018, Belote *et al* 2020, Beyer *et al* 2020), with enormous demand for information about human pressure on the land surface to support policies related to land use change, biodiversity conservation, and climate action (Pascual *et al* 2017, Beyer *et al* 2020, Ruckelshaus *et al* 2020). A key challenge for expanded use of the HFI for operational efforts, however, is that new updates typically lag the present day by seven or more years (Venter *et al* 2016a, Williams *et al* 2020). This time lag means that large-scale or substantial changes to the land surface caused by human activities can occur well before we detect them, hampering our ability to monitor and respond to their effects on biodiversity. This temporal mismatch pertains directly to the achievement of the United Nations Sustainable Development Goals (SDG) since the deadline for their achievement is 2030 (Sachs *et al* 2020b, Naidoo and Fisher 2020, Anon 2020), with some specific targets within SDG15, “Life on Land”, set for even sooner (Sachs *et al* 2020a). While international (Sachs *et al* 2020a) and voluntary national-scale assessments of various biodiversity metrics (Government of the Co-operative Republic of Guyana 2019) are available for monitoring progress toward SDG15, comparing those SDG15 assessments with an operational interpretation of human influence on the terrestrial land surface - one that can be easily and regularly updated - would provide synchronous insights on the social-ecological outcomes and processes of landscape change.

**Figure 1:**
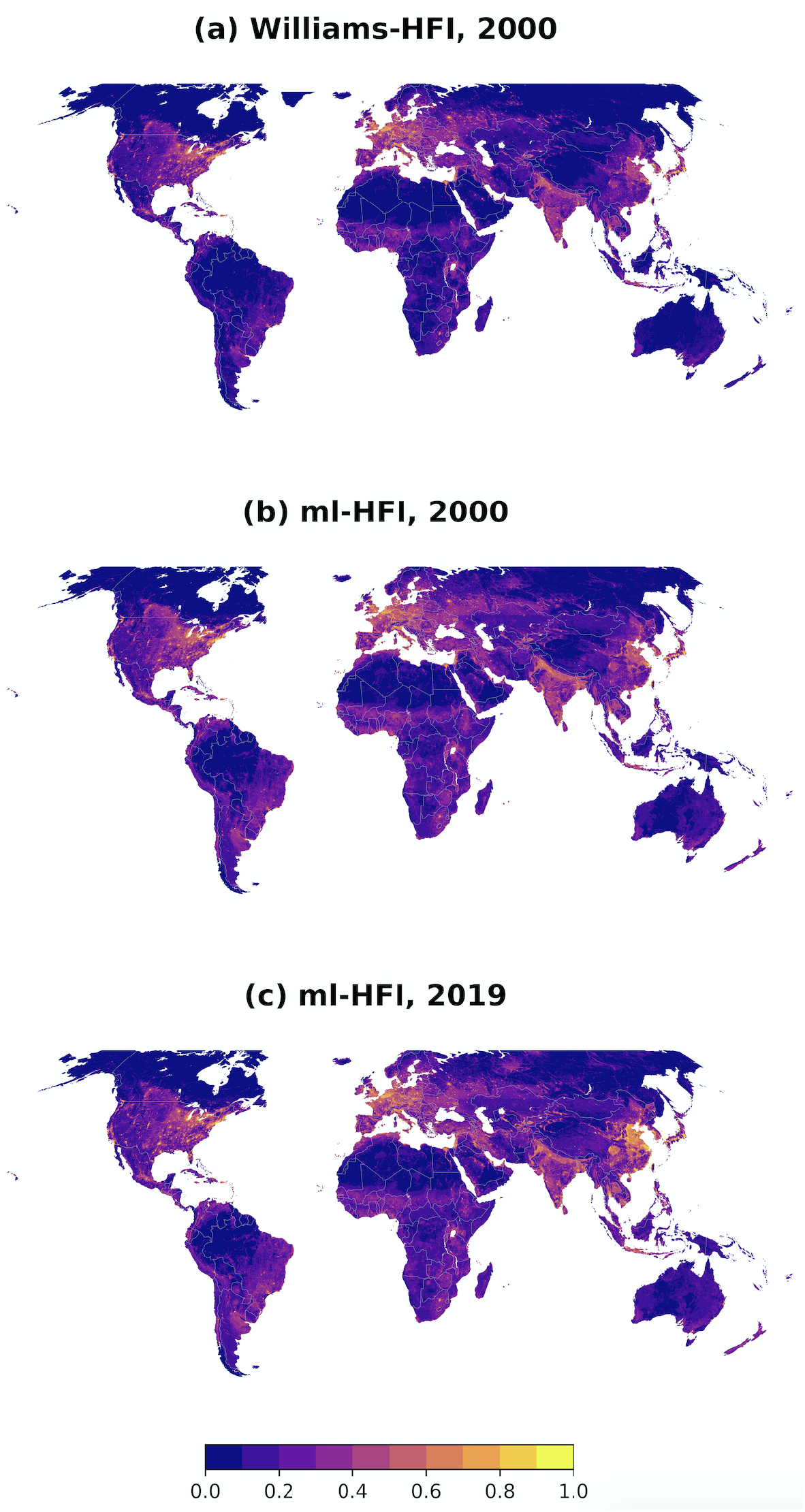
The human footprint index (HFI), a measurement of human pressure on the landscape scaled between 0 and 1. (a) Year-2000 HFI as computed by Williams et al. 2020, (b) year-2000 HFI as computed by a machine learning method, (c) as in (b) but for year-2019.

The conventional approach for creating the HFI (Fig 1a) requires a harmonization of eight different sub-indices representing different aspects of human pressures on the terrestrial surface of the Earth (Williams *et al* 2020). These sub-indices include built infrastructure, population density, and a variety of land use and land cover data. Quantification of the global HFI in this way is a time intensive process and the most recent version of the HFI (published in 2020 (Williams *et al* 2020)) covers years up to 2013.

Machine learning, specifically convolutional neural networks (CNN), are well suited to the task of identifying patterns in imagery (Krizhevsky *et al* 2012, Jean *et al* 2016, Xie *et al* 2015), and have been leveraged extensively to identify various patterns of human activity (Hoffman *et al* 2011, Kumar *et al* 2019, Qiu *et al* 2020). Here, we train a CNN to ingest space-borne imagery of the earth’s surface and predict the human footprint index. Specifically, the Hansen Global Forest Change imagery version 1.7 (GFCv1.7) (Hansen *et al* 2013) includes global, cloud-free, growing season Landsat imagery for the years 2000 and 2019 (Hansen *et al* 2013). We exploit the overlap between the year-2000 Williams-HFI and the year-2000 GFCv1.7 imagery for training the CNN. We then use the trained CNN to predict the 2019 ml-HFI based on 2019 GFCv1.7 imagery. The result is the first near-present, global, machine learning-based human footprint index (ml-HFI) for the years 2000 and 2019 (Fig 1).

While a near-present, global ml-HFI opens many avenues for research, we demonstrate its application to global conservation policy, by comparing it against country-level progress being made toward SDG15. Specifically, using the most up-to-date trends from the United Nations, we find that 43 countries are documented as making progress toward SDG15 and yet also have experienced increases in ml-HFI between 2000 and 2019. Progress toward SDG15 can come about in multiple ways, so we analyze 8 of the 43 countries to visualize how the geographic distribution of changes in human pressure coincide with specific indicators of conservation success. Given that the near-present ml-HFI could be used to motivate specific types of conservation activities, we further explore the policy mechanisms that may support the coexistence of SDG15 progress and increased human footprint. We anticipate future updates to the ml-HFI as soon as updated cloud-free remote sensing imagery is made available, which should enable this research to provide continuing support for biodiversity conservation and sustainable development.

## Data

### Human Footprint Index

We quantify human pressure on the landscape using the human footprint index (HFI), which is a global, dimensionless index of human pressure on the land surface (Sanderson *et al* 2002, Venter *et al* 2016a). We employ the most up-to-date version of the HFI (Williams *et al* 2020), which we denote as “Williams-HFI” hereafter in this text. The Williams-HFI is comprised of eight different sub indices representing different aspects of human pressures to the terrestrial surface of the Earth, including 1) extent of the built environment, 2) population density, 3) electric infrastructure, 4) agricultural lands, 5) pasture lands, 6) roadways, 7) railways, and 8) navigable waterways. Here, we rescale the original index, which ranges from 0 to 50, to a range from 0 to 1 (from low to high human pressure). The Williams-HFI data is output in a gridded format, where each gridpoint is 0.00989273 degrees latitude by 0.00989273 degrees longitude.

### Landsat imagery

We use the Global Forest Change dataset, a global analysis of forest cover change based on Landsat imagery (Hansen *et al* 2013). The data consist of processed Landsat imagery from the bands 3, 4, 5, and 7. The global processing identifies growing-season imagery and only includes cloud-free images. We specifically use bands 3, 5, and 7 in our analysis (corresponding to the “red” band, and two bands of “near infrared”). We chose these bands based on preliminary sensitivity tests, which revealed that these bands provided sufficient spectral specificity for identifying terrestrial changes. The latest version of the Global Forest Change product includes cloud-free processed Landsat imagery for the years 2000 and 2019 (GFCv1.7). However, when cloud-free imagery was not available for the specific year, imagery was taken from the closest year with cloud-free data (for year 2000, data could come from 1999-2012; for 2019 could come from 2010-2015).

### Sustainable Development Goal trends

Data indicating the country-level progress toward Sustainable Development Goal (SDG) 15, ‘Life on Land’ is taken from the latest SDG report (Sachs *et al* 2020a). Broadly, the SDG report serves as the most up-to-data compilation of progress toward all SDGs, drawing from both official sources (e.g.,the United Nations, the World Bank) and from unofficial sources (e.g. research institutions and non-governmental organizations). The three types of data that were used to create the SDG15 trend for each country includes published research on the fraction of protected forest-habitat (Curtis *et al* 2018), fraction of protected freshwater-habitat (Lenzen *et al* 2012, 2013), and threatened “red list” species (Butchart *et al* 2007). The report includes the most current assessment of the trend in SDG15 progress, in terms of ‘on track to achieve the SDG’, ‘moderately improving’, ‘stagnating’, or ‘decreasing’. In this work, we denote ‘on track to achieve the SDG’ and ‘moderately improving’, as both indicative of progress toward SDG15.

### The convolutional neural network method

We train a neural network to take a single Landsat GFCv1.7 image as input and output a single value which represents the network’s prediction of the human pressure on the land surface. For this task, we employ convolutional neural networks (CNNs), a class of neural networks often used in image applications due to their ability to identify specific shapes (or features) in the data that assist in making accurate predictions. We refer the reader to (LeCun *et al* 2015, Géron 2019, Yamashita *et al* 2018) for detailed descriptions of CNNs for image applications of machine learning. The specific CNN design and sampling technique utilized here is outlined in detail in the Supplementary Material for transparency and reproducibility.

The input into the CNN is an array with shape (120,120,3) representing 3 Landsat channels of 120×120 pixels each. A Williams-HFI pixel is approximately 40×40 Landsat pixels. We found that the model performed best when the input included surrounding Landsat pixels which gives the CNN more context to predict the ml-HFI in the center of the image. Thus, we input 3 120×120 pixel Landsat images and task the CNN to output a single value between 0 and 1. This value represents the ml-HFI in the center (see white boxes in Supp. Fig. 2). The CNN is trained to minimize the mean squared error of the predicted ml-HFI compared to the Williams-HFI for the year 2000 when the Williams-HFI and the GFCv1.7 imagery overlap. The model is then frozen (i.e., the CNN weights are fixed) and used to predict the ml-HFI for the year 2019 based on the 2019 GFCv1.7 imagery. In doing this, we make the assumption that the spectral characteristics of individual land-use in the year 2019 are similar to the spectral signatures of the same land-uses in the year 2000, a well-tested method in machine learning-based analyses of land-cover change (Curtis *et al* 2018, Omeiza 2019). Moreover, we have performed manual quality control on 2000 locations globally via Google Earth Pro’s time lapse feature (see Results).

We train a separate CNN for each region shown in Supp. Fig. 1 between 70S and 70N. Different biogeographic areas across the globe provide different challenges (e.g., rocky terrain, deserts) for training the CNN. In addition, the types of human impacts vary across regions as well. We found that training separate CNNs for each region improved the accuracy of the predictions compared to requiring one CNN to describe all possibilities. It is very possible that increasing the amount of training data (see discussion below), and utilizing a more complex CNN architecture, would alleviate this step and allow one to train a single CNN for the entire globe. The CNN trained on Outer Asia was used to evaluate pixels within Oceania due to the small sample size of the region. The model used to evaluate Northern Africa was trained on all of Africa but only evaluated over Northern Africa. A separate CNN was trained over Southern Africa. Pixels used for training are shown in Supp. Fig. 3 (purple shading), and were chosen to optimize computation time while maximizing the representation of unique geophysical features and human impacts.

Over all trained CNNs, we train on 3.8 million Williams-HFI pixels and predict 1.4e8 ml-HFI values for 2000 and 2019, each. That is, we only train on approximately 3% of the non-water data available due to computational and storage limitations, but expect many more samples could be included to further improve the ml-HFI accuracy.

## Results

### Human pressure increasing globally

Global, regional, and local patterns of human footprint from the Williams-HFI (Fig 1a) are identified in the year 2000 ml-HFI with high fidelity (Fig 1b). High accuracy is realized over all levels of human footprint by training on only 3% of the terrestrial locations (Supp Fig 3), with a mean absolute error on unseen testing data of 0.07 (Supp Fig 6). With that said, there are regions of the planet that are systematically more challenging for the CNN to reproduce (Supp Fig 8), including rocky areas of remote deserts and high latitude mountain ranges. Training the model on a larger portion of locations is expected to overcome this limitation in future iterations. Once trained, the CNN is then tasked with predicting the most current version of the ml-HFI from 2019 GFCv1.7 imagery (Fig 1c). This 2019 map represents the most current prediction of the human footprint index (previously most-updated for 2013 (Williams *et al* 2020)).

Changes in ml-HFI between 2000 and 2019 are visualized globally in Fig 2 with positive values indicating a large increase in ml-HFI, and negative values indicating large decreases in the ml-HFI (where “large” is defined as changes greater than 0.25). Patterns of change are consistent with a steady expansion of human encroachment into low ml-HFI areas, as well as increasing density of pressure in areas of higher ml-HFI. Inset panels in Fig 2 display the GFCv1.7 imagery from 2000 (left panels) and 2019 (middle panels) for different forms of increasing human pressure.

**Figure 2:**
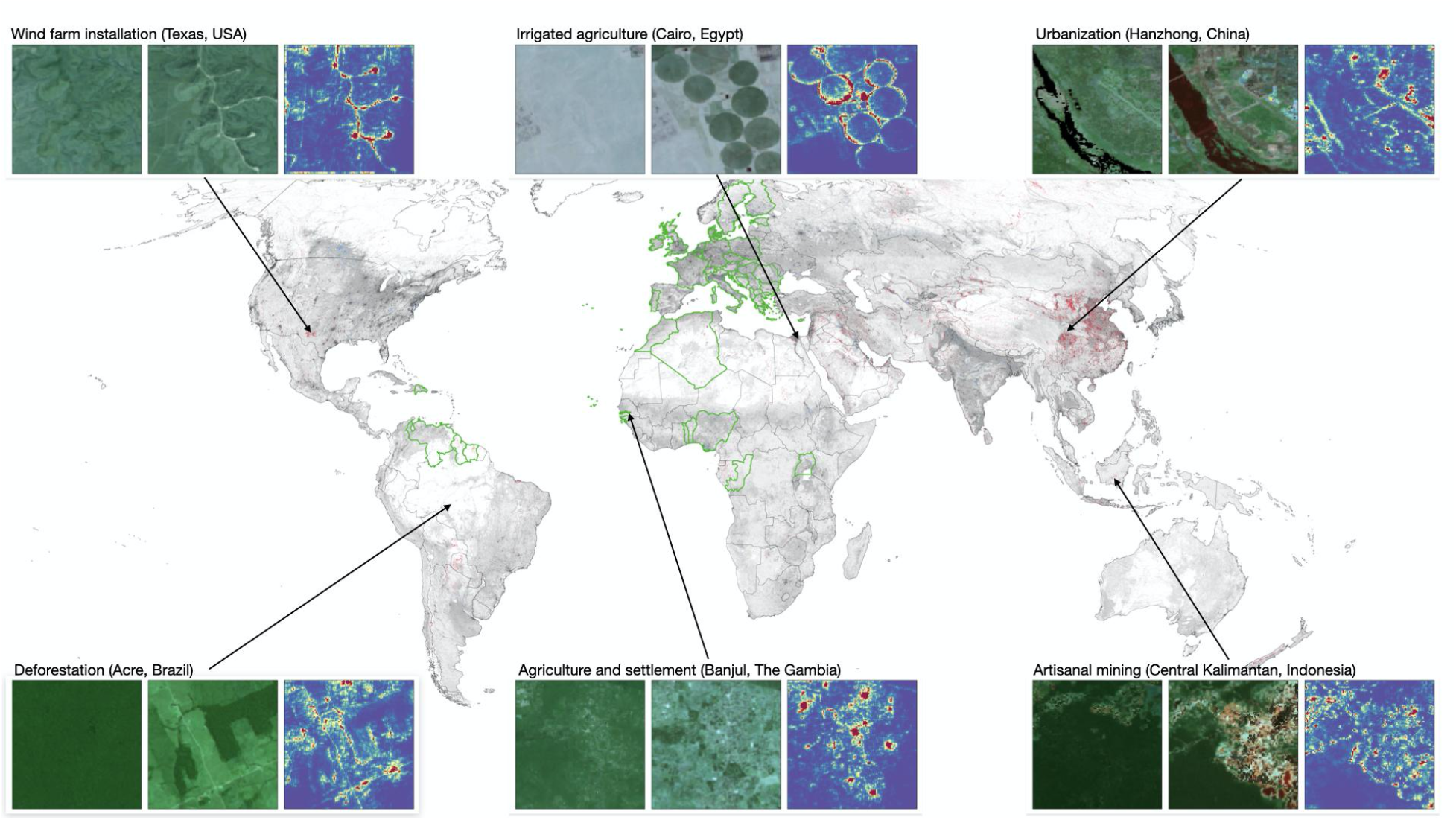
Large changes in human pressure defined as the ml-HFI in 2019 minus that in 2000, where red shading denotes changes larger than 0.25 and blue shading denotes changes smaller than −0.25. Gray shading denotes the ml-HFI from 2000 for reference. Green outlined countries are experiencing substantial increases in HFI and making progress toward SDG15. Inset panels provide examples of increasing human pressure and the relevant features used by the CNN to identify human activity. From left to right, each inset shows (left) year-2000 GFCv1.7 imagery, (middle) year-2019 GFCv1.7 imagery and (right) features most relevant to the CNN for its year-2019 prediction of the ml-HFI. The GFCv1.7 imagery is plotted in false color as its spectral bands are outside of the visible spectrum.

### Attribution of the CNN’s Decisions using Layer-wise Relevance Propagation

While the human eye can easily see the changes in the images from one year to the next, we leverage a neural network visualization technique called Layerwise Relevance Propagation (LRP) (Bach *et al* 2015, Montavon *et al* 2017) to reveal the features that the neural network deems most relevant for its calculation of the ml-HFI (right panels). This method highlights the regions of the input, or the pixels of the 3 Landsat images, that were most relevant to the CNNs output (i.e., prediction of the ml-HFI). This technique acts to “open the black box” for neural network tasks and, importantly, provides intuitive confirmation or possible refutation of the CNN’s decision-making process. In our case, we use LRP as a quality control step to check that the CNN is paying attention to the correct features that would lead to high or low ml-HFI predicted values.

LRP, applied separately to each prediction of the CNN, produces a heatmap displaying the relevance of each pixel of each of the 3 input Landsat images (i.e., channels). We have computed the LRP relevance heatmaps for selected predictions of the 2019 ml-HFI where human pressure has substantially increased. The average relevance across the 3 input channels are displayed in Fig. 2 and in Supp. Fig. 2 where warmer colors denote pixels that were most relevant to the ml-HFI prediction of the CNN. For all of these examples, the CNN places the highest relevance on features within the image that are clearly due to human activity as seen by eye in the 2019 Landsat imagery. This provides us with confidence that the CNN predictions are correct for the right reasons.

### Evaluation of CNN results

We assess the performance of the year-2000 ml-HFI relative to the year-2000 Williams-HFI by examining both the accuracy (Supp. Fig. 5) and absolute error (Supp. Fig. 6). The ml-HFI values are within an error tolerance of 0.1 of the Williams-HFI values for 73% of the terrestrial globe. When the error tolerance is increased to 0.2, the percentage jumps to 93%. Mean absolute errors as a function of the Williams-HFI values (binned in increments of 0.1) are shown in Supp. Fig. 6.

The median absolute error is well below 0.1 for most bins, and this is especially true for regions of very low or very high ml-HFI. The training samples and testing samples generate similar error distributions, which suggests that the CNN models were properly trained. Global maps of the errors and absolute errors are displayed in Supp. Fig. 7-8.

To demonstrate the success of the ml-HFI in predicting human pressure from GFCv1.7 Landsat imagery, Supp. Figs. 9-12 provide example outputs of the ml-HFI for urban, agricultural, mining, and rural landscapes in the year-2000. In all of these cases, the ml-HFI compares well with the Williams-HFI, although it is clear that the Williams-HFI emphasizes road networks and is smoother than the ml-HFI which is predicted on a pixel-by-pixel basis.

We assess the skill in the ml-HFI values using the Williams-HFI values as “truth”; however, the Williams-HFI values themselves are not ground truth, but are estimations that can also be incorrect at times (although this is likely rare due to the extensive amount of work that goes into developing this index). With that said, we have found multiple locations where the ml-HFI predicts high human pressure but the Williams-HFI does not. Upon inspection of these cases we believe the ml-HFI to be correct. Four such examples, provided in Supp. Fig. 13, highlight how the ml-HFI can be useful for identifying regions of high human pressure that may have been overlooked in the Williams-HFI.

There are also regions where we believe the ml-HFI struggles. The largest errors appear over dry, rocky terrain devoid of human activity, for example, the rocky parts of the Saharan Desert, the Australian desert, and the spine of the Andes (Supp. Fig. 7-8). In these areas, the ml-HFI predicts human activity where none exists. We provide three such examples in Supp. Fig. 14. However, ml-HFI errors over these problematic regions with low Williams-HFI are still relatively small (Supp. Fig. 6-8), with the 90th percentile error being approximately 0.12. Because the mI-HFI appears to struggle over rocky deserts and mountains where the Williams-HFI is zero, we mask out regions where the Williams-HFI was less than 0.02 in 2000 (termed “Wilderness” by (Williams *et al* 2020)) and the mI-HFI error was larger than 0.001 for our change calculations. That is, we mask out changes where the 2000 mI-HFI initially struggled with a zero prediction (i.e. no human activity at all).

### Biodiversity gains despite development

Previous research underscores the transformative effects that human activities have on natural ecosystems (Biggs *et al* 2009, Rocha *et al* 2018). Many of these studies indicate that heightened human pressure on the landscape diminishes the capacity for those lands to support biodiversity. We interrogate those assumptions using the ml-HFI for the years 2000 and 2019 with a contemporary assessment of country-level progress toward achievement of SDG15 (Sachs *et al* 2020a). Globally, there are 119 countries experiencing increases in the ml-HFI. We find that 76 of these countries are experiencing either stagnating or decreasing progress toward SDG15 (not shown). This supports the common belief that an increasing human footprint is not compatible with the maintenance or improvement of terrestrial biodiversity. Contrary to this common belief, however, we also find that 43 countries (Fig 2, green outlines) are experiencing substantial increases in human pressure over a large fraction of their land surface (increases of 0.25 or larger over 1% or more of their area) while at the same time “moderately improving” or actively “on track to achieve” SDG15. In other words, 36% of countries experiencing increases in human pressure, are also making progress toward SDG15.

These substantial changes identified by the ml-HFI have been further confirmed manually for more than 2,000 locations across the 43 countries using the time lapse feature in Google Earth Pro. Norway, Gabon, Libya and Sudan have been removed from the list as further inspection shows that the CNN struggles in these locations. In Norway the presence of high latitude rocky areas are mis-classified as experiencing change. In Gabon there is an apparent GFCv1.7 imagery issue which appears to be localized specifically to Gabon’s interior. Both Libya and Sudan contain large areas of rocky, remote deserts that are misclassified as experiencing substantial human impact.

Our core finding, that the ml-HFI is increasing while at the same time biodiversity gains in some countries are also increasing, remains nascent in the conservation and development discourse (Sarkar 1999, Newbold *et al* 2016, Ellis *et al* 2012, Watson *et al* 2018). To better understand this apparent coexistence, we summarize the types of ml-HFI changes across a subset of eight countries along with specific SDG15 indicator progress in Fig 3. The ml-HFI reveals a wide spectrum of human pressure on the landscape, including: deforestation for extensive mining (Suriname, Guyana), spreading road networks (Estonia, Slovenia), urbanization (Uganda, The Gambia), and expanding agriculture (Benin, Morocco). Moreover, there is no uniform pathway for making progress across the SDG15 indicators for forest-(Curtis *et al* 2018), freshwater- (Lenzen *et al* 2012, 2013), and threatened “red list” species conservation (Butchart *et al* 2007). For example, Slovenia and Estonia are on track to achieve SDG15 in all three indicators. But for most countries, it is a very mixed pathway across the three indicators. For example, Uganda may be making progress on forest and freshwater conservation, while decreasing toward its “red list” species target. Evidently, the types of increasing pressures revealed by the ml-HFI do not lead to identical patterns of SDG15 progress. This suggests additional country-level policy information may deepen our understanding and we provide three such case studies: Guyana, The Gambia, and Morocco.

**Figure 3:**
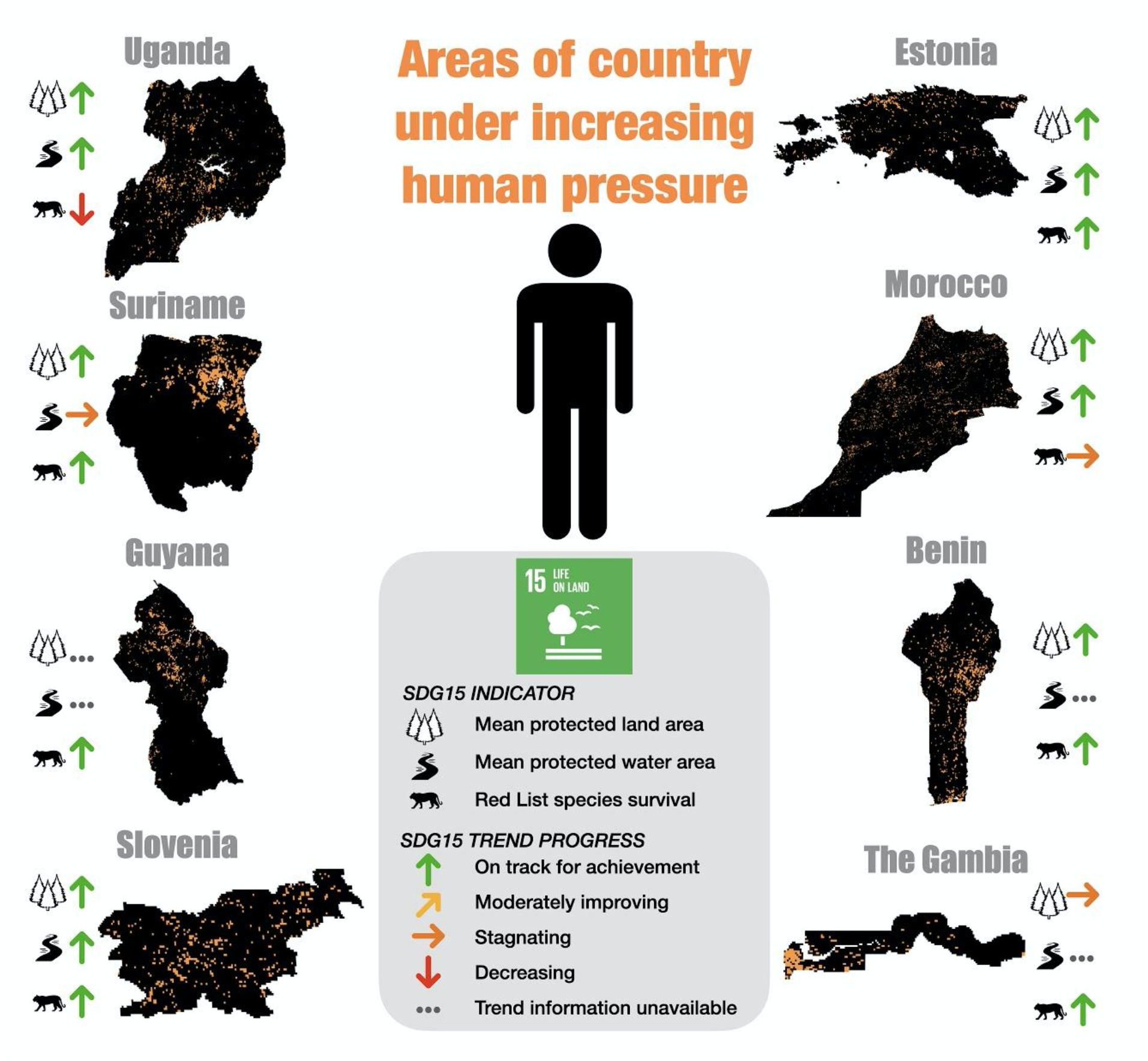
Countries experiencing substantial changes in human pressure that are also making progress toward SDG15. Percent indicates the fraction of each countries’ land surface area that is experiencing substantial increases in human pressure, defined as a change of 0.25 on the 0-1 scale of the ml-HFI, over land surface area greater than or equal to 1% of the total country area. Countries shown are a subset of the full list of 43 countries meeting these criteria detailed in Supp Table 1. ml-HFI changes displayed here are degraded in resolution for easier visualization.

### Cases of development processes

A core function of the ml-HFI is to monitor remote regions experiencing rapid change, as clearly identified in Guyana - including considerable expansion of road networks, deforestation and extensive mining in the forested interior (Fig 3, Supp. Fig. 15). Notwithstanding its accelerating economic development, Guyana has the second-highest per capita forest cover in the world, second only to its neighbor, Suriname (Government of the Co-operative Republic of Guyana 2019). Given the enormous potential for forest preservation, Guyana was one of the first countries to implement REDD+ (Reduced Emissions from Deforestation and Forest Degradation), partnering with Norway which provides financial resources to incentivize the reduction in deforestation rates, and the establishment of protected areas in collaboration with indigenous communities (Government of the Co-operative Republic of Guyana 2019). These activities contribute to Guyana’s current progress toward its SDG15 goals and, if continued, will lead to consistently improving trends in forest and freshwater habitat. Yet, the growing network of roads and mining across the Guiana Shield revealed by the ml-HFI shows how quickly development processes can encroach into wild and remote areas, and may threaten the continued success of biodiversity conservation efforts.

Urbanization of agricultural areas is a key feature of development (Ellis and Ramankutty 2008), and as evidenced by the ml-HFI results for The Gambia, can take place explosively over the course of less than 20 years (Fig 3, Supp. Fig. 16). Elsewhere in The Gambia, the ml-HFI reveals changes driven by deforestation from roads and agriculture. The Gambia’s Voluntary National Review for SDG15 highlights the role that climate change has had on exacerbating challenges to natural resource management, as well as the tension between preserving biodiversity with food security for its population - which remains 50% rural (The Republic of Gambia 2020). Nonetheless, The Gambia includes multiple protected forests, including biodiverse areas near the mouth of the The Gambia River. The ml-HFI results for 2019 show the lowest levels of human pressure in precisely these riparian locations, emphasizing the need for continued protection of these forests.

Inspection of the ml-HFI shows that Morocco is experiencing development pressure in the form of expanding peri-urban farmland, as well as road and energy infrastructure (Fig 3, Supp. Fig. 17). As a middle income country, with both semi-arid forests and desert ecosystems, Morocco has focused on boundary demarcation of forested areas, enforcement of forest protection from illegal activities, and expanding coverage of forest management plans (Royaume du Maroc 2020). Such efforts are representative of Morocco’s capacity for management and enforcement, and suggest that advanced monitoring tools, like the ml-HFI, could contribute important and actionable information. As with Guyana and The Gambia, development is still taking place and, as clarified in Morocco’s Voluntary National Review for the SDGs, there are ongoing plans for water and land development as well as forest conservation in Morocco (Royaume du Maroc 2020). The ml-HFI results for 2019 support the assessment that considerable potential exists for conservation, especially in the remote mountains and arid ecosystems of the east and southern reaches of Morocco.

### Prospects for development monitoring

Combining the SDG15 progress with the 2019 present-day human footprint index shows that progress toward biodiversity goals can be made despite increases in human pressure. These increases occur in countries ranging in wealth, economic development, or reliance on specific natural resources. For many countries that are aiming for resource intensive economic development, monitoring and evaluation will be critical for tracking changes in near-real time.

Countries that are still actively developing may find the ml-HFI useful to identify strategies to provide effective guardrails for staying on track for protecting biodiversity (e.g., Dominican Republic, Republic of Congo), especially when increased human activity encroaches on pockets of biodiversity. For wealthier or more developed countries with longstanding regulatory structures for protected areas, the ml-HFI can help determine conservation policy efficacy and ensure that additional increases in ml-HFI concentrate in already developed areas (e.g., Sweden, Switzerland).

## Conclusions

Although we detail compelling evidence of progress toward biodiversity in the face of increasing human pressure for some countries, the more common pattern is that of increasing human pressure at the expense of gains in biodiversity conservation. We do not single out any specific countries that are not on track to achieve SDG15 (Sachs *et al* 2020a), but the changes in their ml-HFI results reveal similar pressures to the cases detailed above - urban growth, agricultural expansion, and resource extraction (mining and timber harvesting). While considerable efforts are being made to protect intact ecosystems from encroaching or intensifying human activities, growth in human footprint is likely outpacing field-based monitoring protocols to assess environmental impacts and highlight that many current policies are insufficient to regulate development (Curtis *et al* 2018). Capable of capturing near present changes to the land surface, the ml-HFI could be an important tool for assisting countries that are flagging behind their SDG15 targets to close the gap by 2030. We also acknowledge that this is version 1.0 of the ml-HFI, and that with greater computational power, for instance, more data could be used for training, with corresponding increases in accuracy. Furthermore, the use of innovative tools such as Layerwise Relevance Propagation indicates that the machine learning-based approach is correctly identifying human-made features and activities (e.g., transportation networks or landcover conversion) providing confidence in, and interpretability of, the CNN’s predictions. This step is especially important if the ml-HFI is to be used for policy decisions as national and international laws have begun to require explainable AI systems for decision-making (High Level Expert Group on AI 2019, Phillips *et al* 2020).

That some countries can simultaneously be progressing toward sustainable development goals while experiencing dramatic increases in human pressure confounds expectations. The near-present, global ml-HFI provides an analytical tool for interpreting this unexpected coexistence. Moving forward, science and policy must work to understand why and how this is possible. Such activities will permit the development of better policies that will foster a thriving society and environment in the Anthropocene.

## Acknowledgments

This research was funded, in part, by the United States National Aeronautics and Space Administration (NASA) under grant #18-SLSCVC18-0006.

## Author contributions

All authors devised the research plan. E.A.B. undertook the development of the convolutional neural network, including design, training and validation. P.W.K. processed sustainable development goal data and performed quality control on neural network model output. All authors contributed to the writing of the manuscript.

## Data Availability and Code Availability

The ml-HFI for 2000 and 2019 as well as the weights for the trained convolutional neural networks will be made available to the community via the Mountain Scholar permanent data repository with a permanent DOI and via github. All data used in this study is publically available. Human Footprint Index data were downloaded from the github repository https://github.com/scabecks/humanfootprint_2000-2013. Sustainable Development Goal data were downloaded from the United Nations website https://dashboards.sdgindex.org/downloads. Landsat imagery was downloaded from the Hansen Global Forest Change v1.7 repository https://earthenginepartners.appspot.com/science-2013-global-forest/download_v1.7.html.

