## Supplementary Materials for "A machine-learning approach to human footprint index estimation with applications to sustainable development" (from another version of this preprint: version 2, 2020-11-02)

**Affiliations:**
[1]School of Global Environmental Sustainability, Colorado State University, Fort Collins, CO
[2]Department of Atmospheric Science, Colorado State University, Fort Collins, CO
[3]School for Environment and Sustainability, University of Michigan, Ann Arbor, MI

**Contents:**

- Supplementary text on the convolutional neural network method
- Supplementary Figures S1-S17
- Supplementary Table S1

### Supp. Text: Additional Material about the Convolutional Neural Network Model

#### Network architecture

The CNN architecture and parameter values were determined through systematic testing to find a set of values that performed well for this data set. Specifically, each CNN takes as input a 2D (latitude by longitude) Landsat image with 3 channels, each channel associated with one of the three spectral bands discussed above. The CNN then consists of 5 convolutional layers with stride of 2 and "zero padding". The first convolutional layer has 64 filters, while all subsequent convolutional layers have 128 and all filters have size 3x3. The first 4 convolutional layers are each followed by max pooling layers (i.e. upsampling layers) with pooling size 2x2, stride of 2, and "valid padding". The output of these layers is fed into a final hidden dense layer of size 32 units, which then feeds into the output layer (i.e. the prediction layer) of size 1 which corresponds to the single predicted ml-HFI value. All activation functions throughout the network use the rectified linear unit (ReLu) except for the dense layer, which utilizes a sigmoid activation function to constrain the final ml-HFI output between 0 and 1.

To try and reduce overfitting during the training of the CNN, we include a "dropout layer" (Srivastava et al. 2014) with a 50% dropout rate following the final convolutional layer but prior to the dense layer. The dropout layer acts as a regularizer by randomly removing 50% of its hidden units during training. This practice serves to reduce overfitting and improve generalization to data not seen by the network during training.

#### Input data

The input into the CNN is an array with shape (120,120,3) representing 3 Landsat channels of 120 pixels x 120 pixels each. As a preprocessing step, the three Landsat input channels (band 3, band 5, band 7) are standardized, first dividing by 255 (to convert to values between 0 and 1), then subtracting (0.08,0.24,0.12) and dividing by (0.46,0.58,0.47). In this way, we obtain values that are centered around zero and extend to approximately plus/minus 1.0. Throughout our analysis, we noticed that the general "brightness" of the Landsat imagery can vary in space, and also between year 2000 and 2019 within the GFCv1.7 data. We add a random integer between (-10,10) to each input channel prior to training to eliminate the tendency of the CNN to base its predictions too heavily on the magnitude of the input values. This step may be exceptionally important if one wishes to train and predict from various versions of the GFC Landsat imagery.

#### Training and loss function

Because across the globe there are substantially more low Williams-HFI pixels than high pixels (see Fig. 1 and Supp. Fig. 4), we restrict training to a maximum of 40,000 pixels per 10 deg. by 10 deg. box, and enforce that no ml-HFI bin is represented by more than about 4,000 samples each. Since high Williams-HFI values are quite rare (see Supp. Fig. 4), we take a standard machine learning approach and perform data augmentation (Shorten and Khoshgoftaar 2019) prior to training to increase the sample size of these rare inputs. Specifically, we augment our training set by rotating each high Williams-HFI input image by 90 degrees to create a new input image (a rotated version of the original). This is done two more times to obtain four different rotated versions of the same satellite image. From the perspective of the CNN, this creates 4 unique training/output samples for a single Williams-HFI pixel from which the CNN can learn.

The CNN outputs a single, continuous variable and is trained to minimize the mean squared error of the predicted ml-HFI compared to the Williams-HFI. This training is performed for the year 2000 data when the Williams-HFI and the Landsat imagery overlap. Initially, we noticed that the trained CNNs tended to avoid predicting the value of zero. A Williams-HFI value of zero holds special meaning, as it implies a pristine landscape with no human influence, and so we modified the loss function to place more weight in getting the zeros correct. This is accomplished by re-labelling the samples with labels of zero to having labels of -0.2 for computation of the loss function during training. Because the sigmoid activation function (which bounds its outputs between 0 and 1) is applied prior to the output of the CNN, the CNN is unable to ever predict a value below 0; thus, this relabeling encourages the CNN to predict values of exactly zero.

We train each CNN using a learning rate of 0.001 and a batch size of 128. The learning rate sets the size of the step one takes in the down-gradient direction of the loss function, while the batch size sets the number of samples to process before evaluating the gradient. After selecting a maximum of 40,000 samples for each 10 deg. by 10 deg. Landsat granule, we withhold 750 samples from that granule for validation during training. The rest of the samples in that granule are used for testing. The 750 validation samples are chosen to ensure that the range of HFI values are all represented at a similar frequency. Training is halted when the validation mean squared error increases for three epochs in a row (i.e. for three complete iterations through all of the samples). At that point, we restore the weights from the best model from prior epochs to define the final CNN.

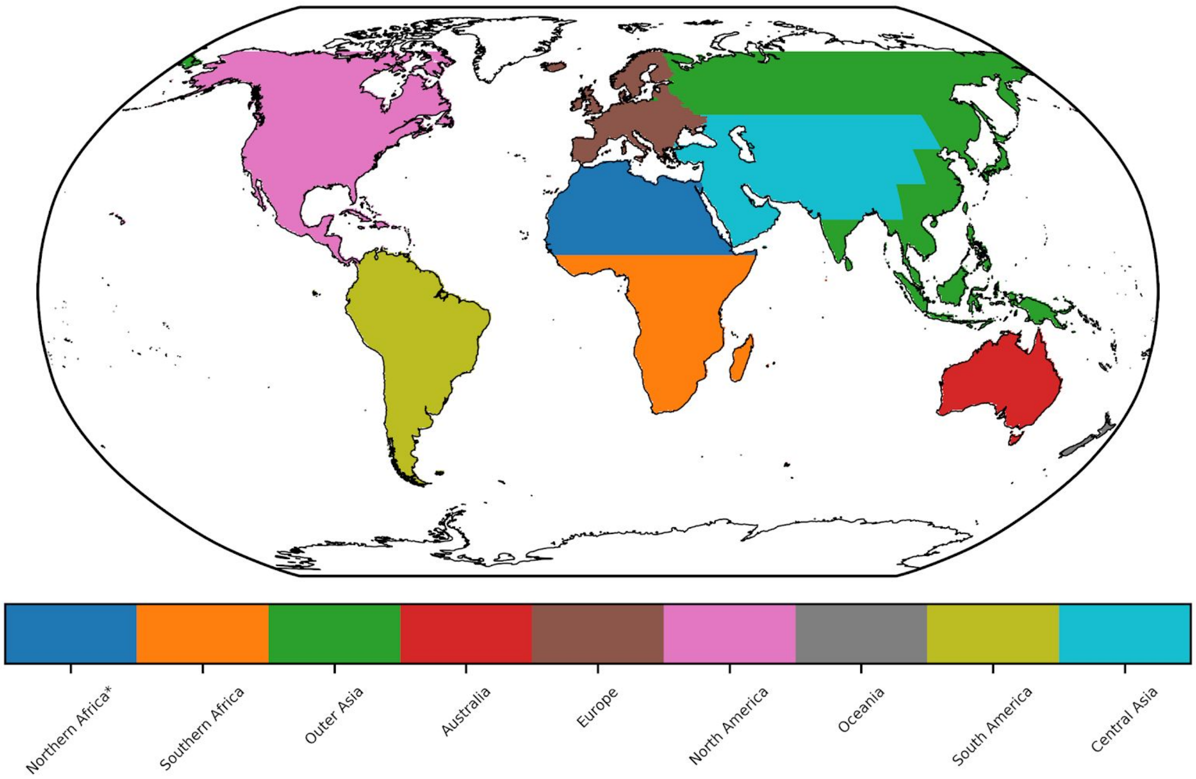


**Supp. Figure 1:** Continental regions for which different CNNs were trained and evaluated. The two exceptions are that (1) the Northern Africa CNN was trained over all of Africa, but only evaluated on Northern Africa samples and (2) Oceania was not used to train a CNN, but instead was evaluated using the Outer Asia CNN.

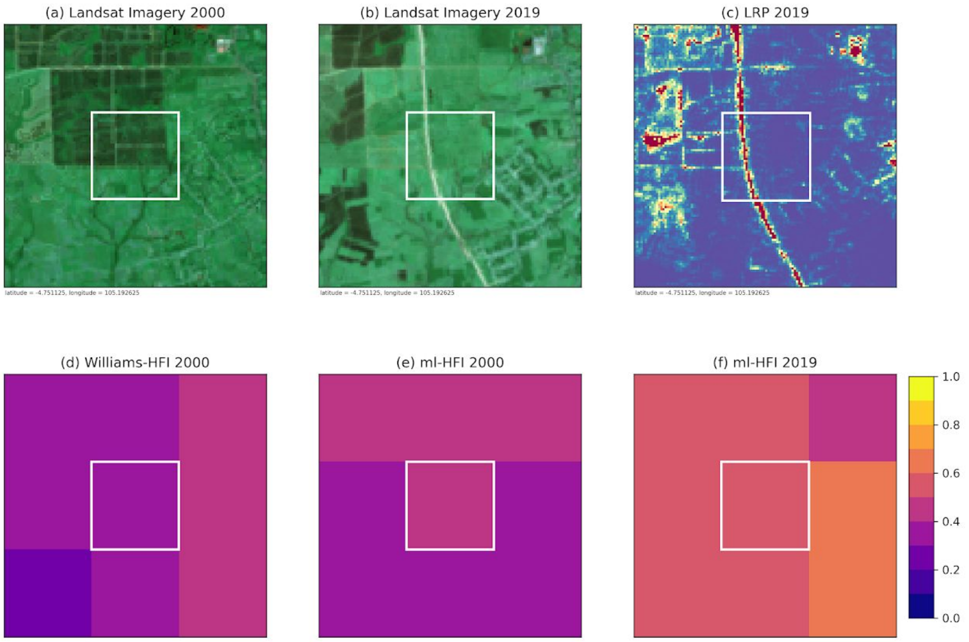


**Supp. Figure 2:** Sample showing the input and output of the ml-HFI and comparison to the Williams-HFI for a region over Sumatra, Indonesia. (a) GFCv1.7 imagery from 2000, (b) GFCv1.7 imagery from 2019, (d) year-2000 HFI from the Williams-HFI, (e) year-2000 HFI from the ml-HFI and (f) year-2019 HFI from the ml-HFI. (c) Displays the heatmap from applying layerwise-relevance propagation (LRP) to the ml-HFI prediction for year-2019. Warm colors denote regions that were most relevant for the CNN's prediction of high HFI. White boxes denote the center 40x40 pixel Landsat box which corresponds to a single HFI pixel for which the ml-HFI predicts the human pressure.

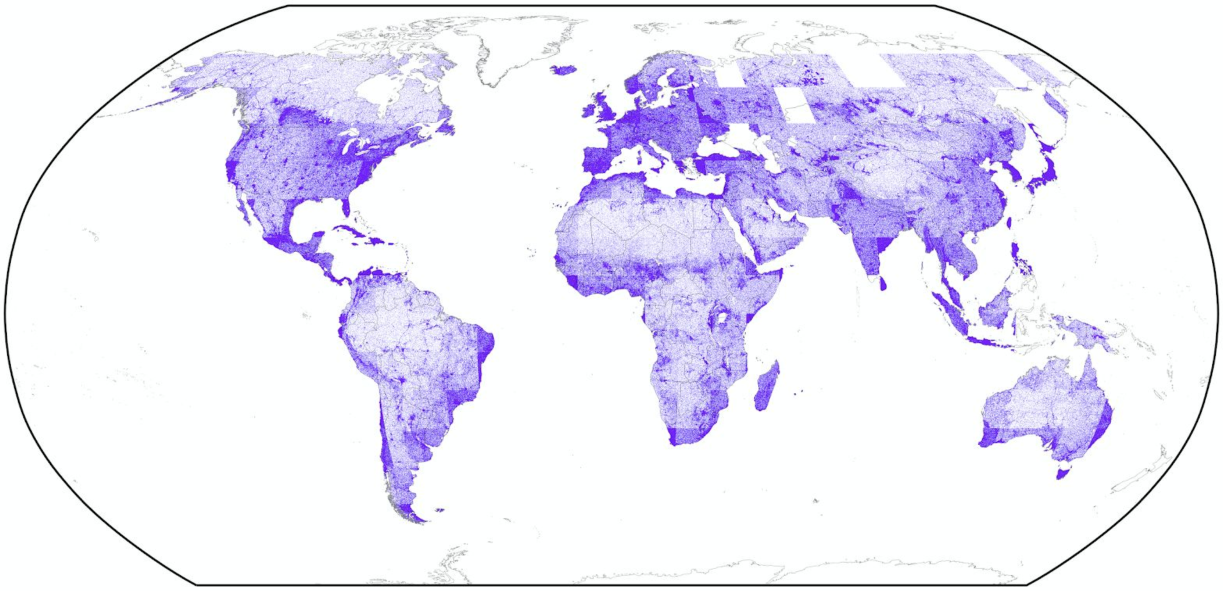

**Supp. Figure 3:** Locations used for training the CNNs. In order to make the image file size manageable, we color an image pixel purple if at least one of the nine HFI pixels within it was used for training. Thus, while it appears that a large fraction of the globe was used for training, in fact only 3% of non-water pixels were used. Some Landsat granules in Outer Asia (refer to Supp. Fig. 1) were not used for training to optimize the time it took to train the CNN for this large region.

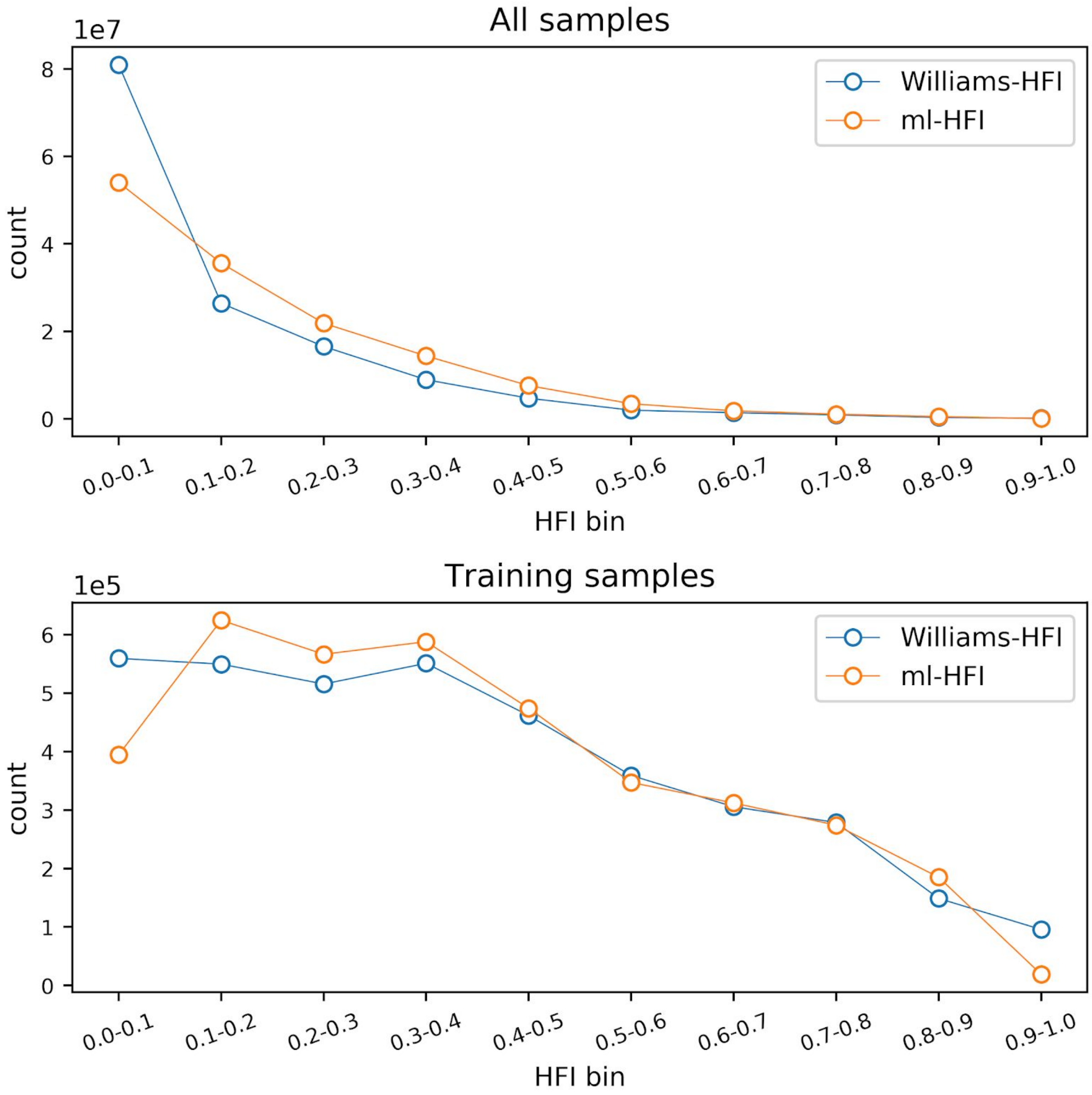


**Supp. Figure 4:** The number of samples per HFI bin (top) across all samples and (bottom) for training samples. Note that the y-axis of the top panel is a factor of 100 larger than that of the bottom panel.

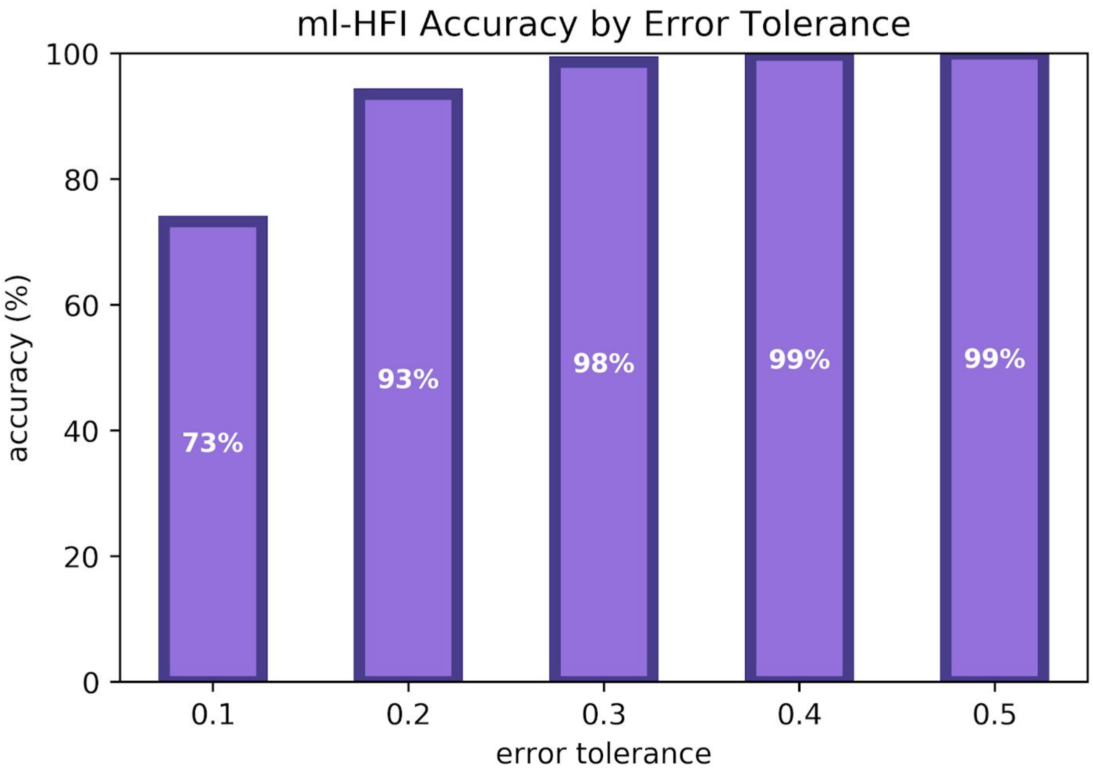


**Supp. Figure 5:** Percent accuracy computed globally for the year-2000 ml-HFI samples as a function of error tolerance. Error tolerance is defined as the maximum allowed absolute error to be counted as a correct prediction. All training and testing samples were included in this calculation.

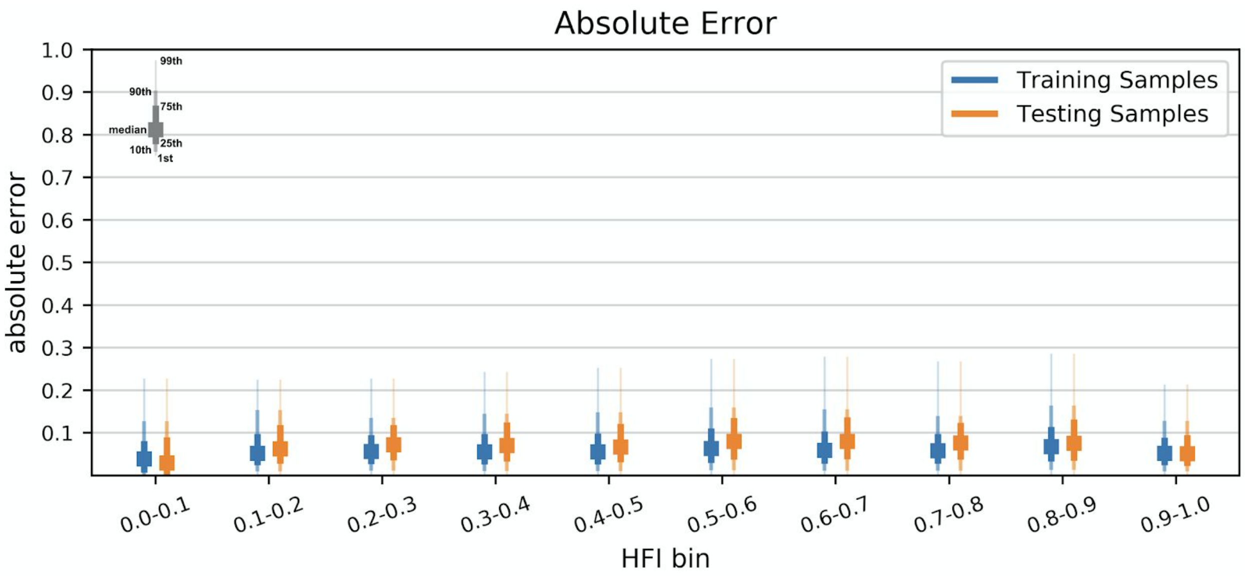


**Supp. Figure 6:** Distribution of absolute errors across all training and testing samples as a function of HFI bin (as defined by the Williams-HFI). The total mean absolute error over all samples is 0.07. Box plots denote the median and a range of percentiles as denoted in the upper-left-hand corner of the plot.

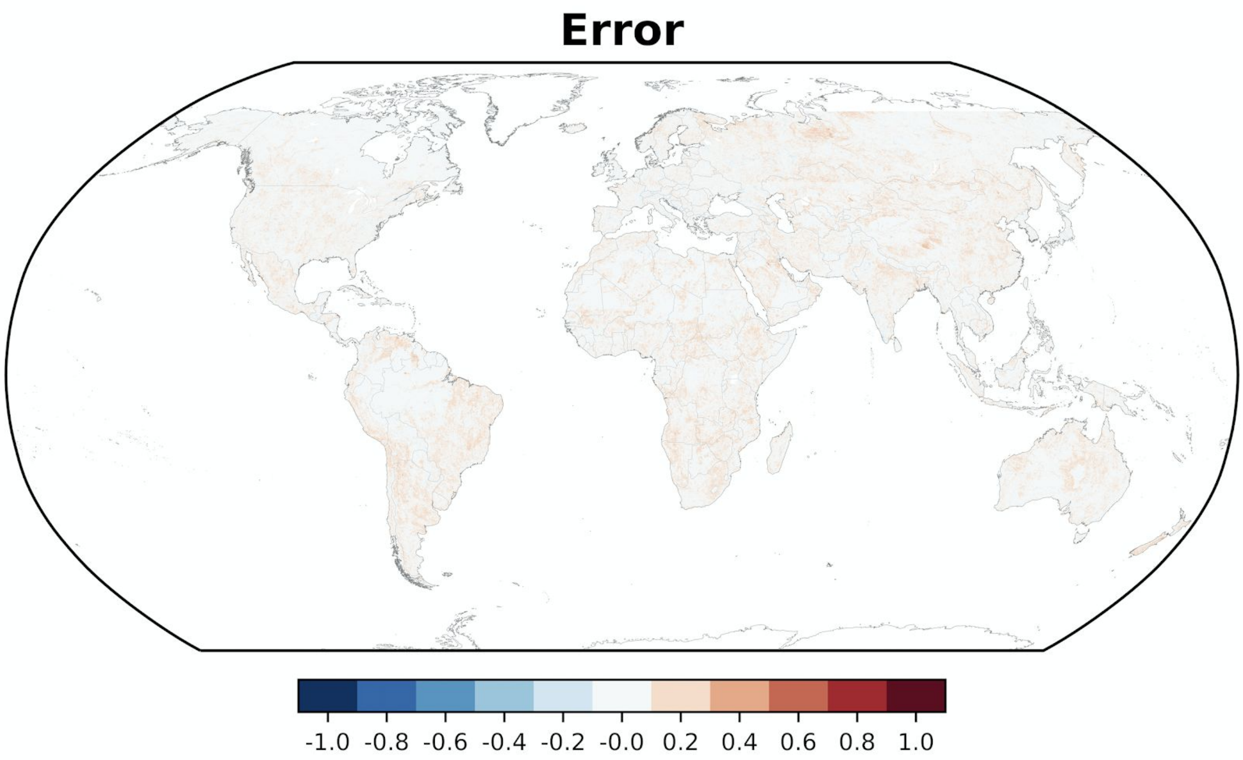


**Supp. Figure 7:** ml-HFI errors defined as the ml-HFI minus the Williams-HFI. In order to make the image file size manageable, the shading within an image pixel is the average error over the nine HFI pixels within it. Note that errors are almost never above 0.3 which is why the colors shown only span a subset of the colorbar range.

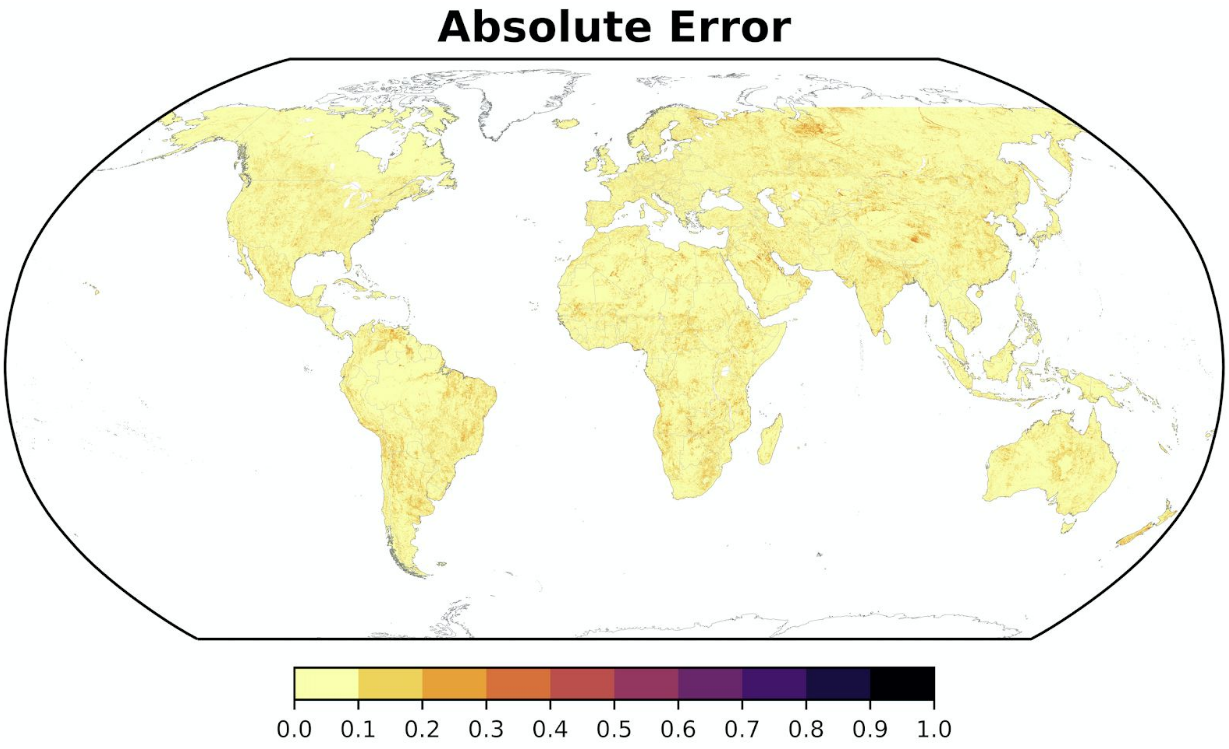


**Supp. Figure 8:** ml-HFI absolute errors defined as the absolute value of ml-HFI minus the Williams-HFI. In order to make the image file size manageable, the shading within an image pixel is the average absolute error over the nine HFI pixels within it.

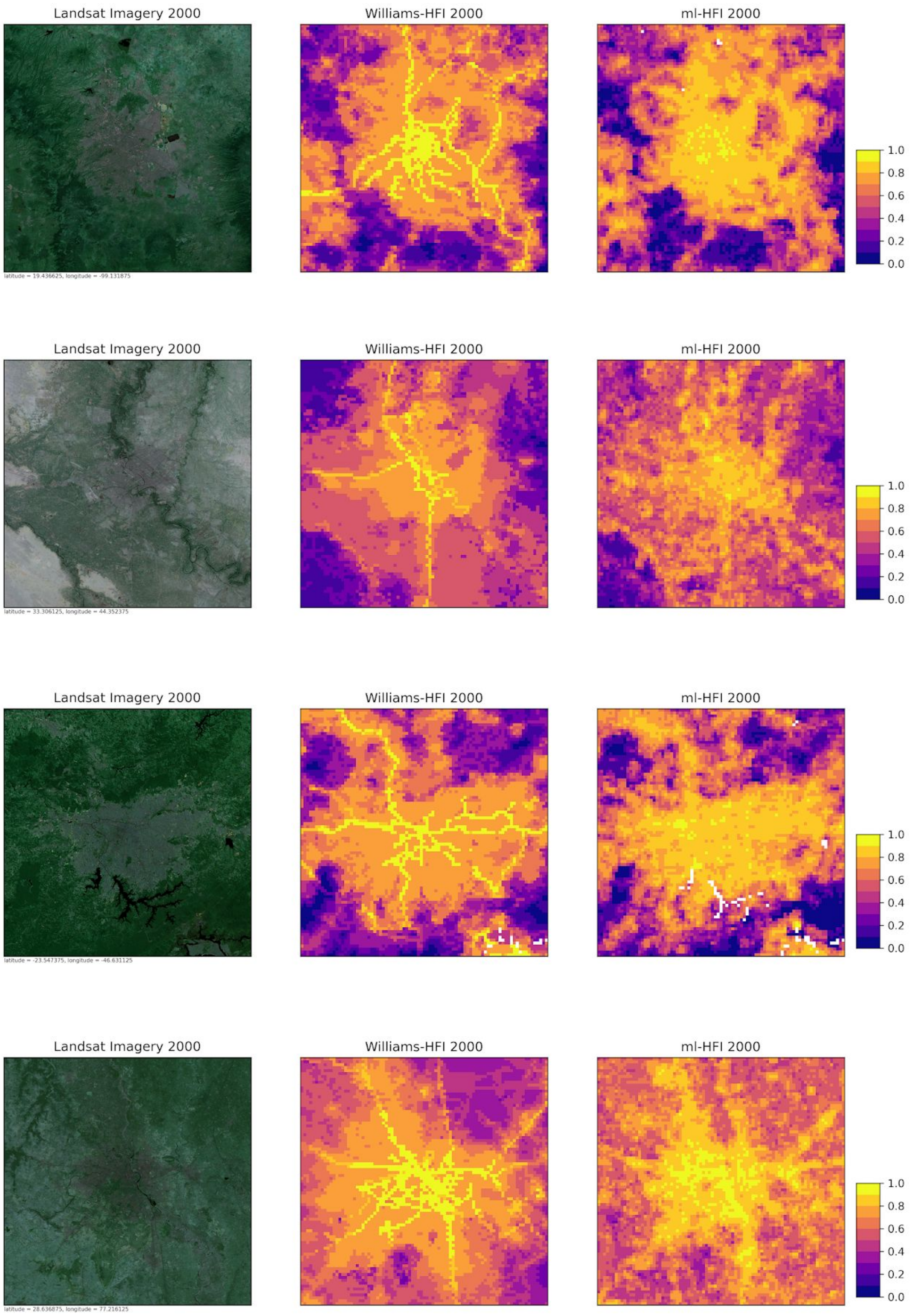


**Supp. Figure 9:** Examples of urban areas where the ml-HFI correctly predicts the human pressure on the landscape. Metropolitan areas highlighted include Mexico City, Mexico; Baghdad, Iraq; São Paulo, Brazil; New Delhi, India from top to bottom.

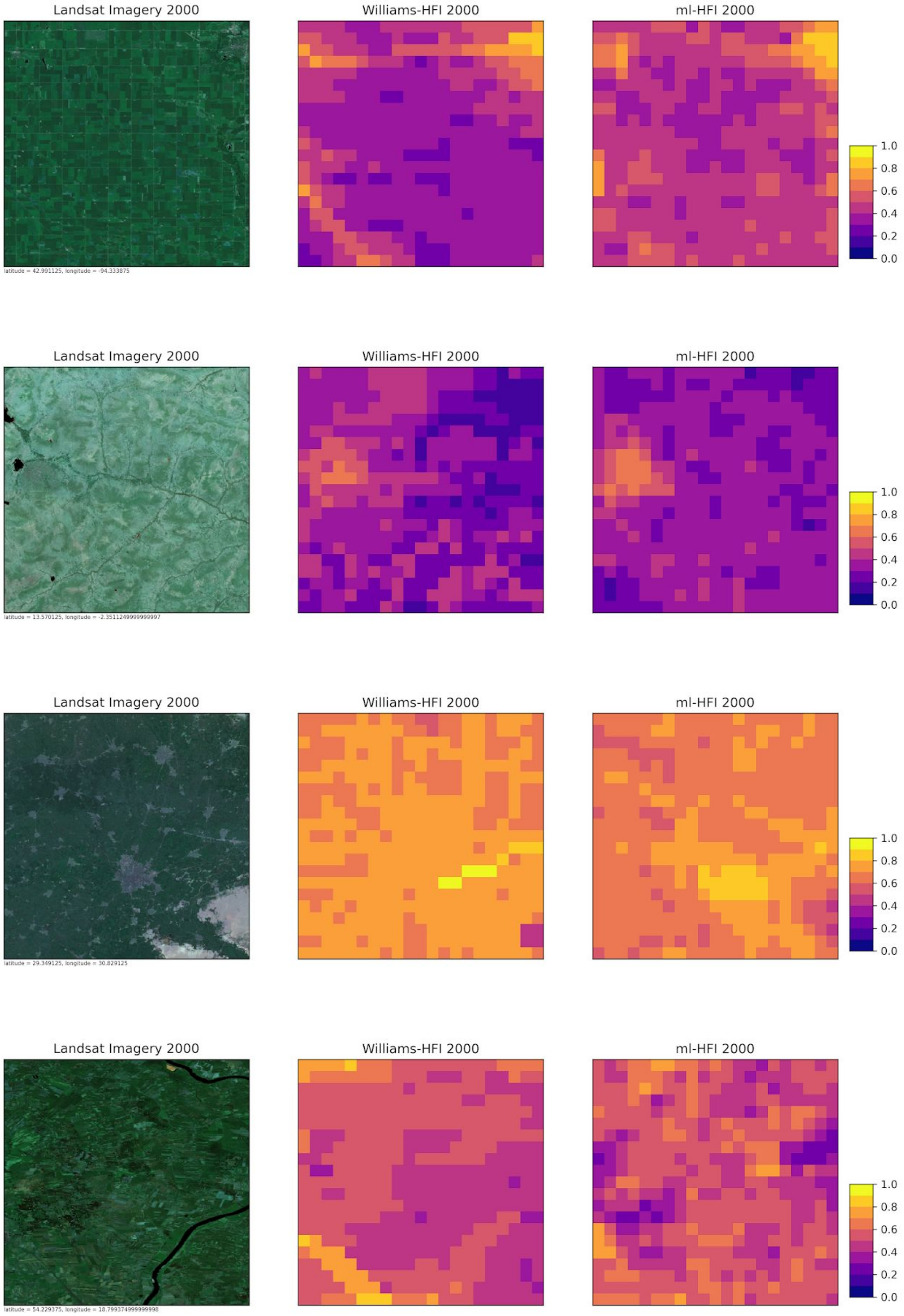


**Supp. Figure 10:** Examples of agricultural areas where the ml-HFI correctly predicts the human pressure on the landscape. Regions highlighted include USA; Burkina Faso; Egypt; Poland from top to bottom.

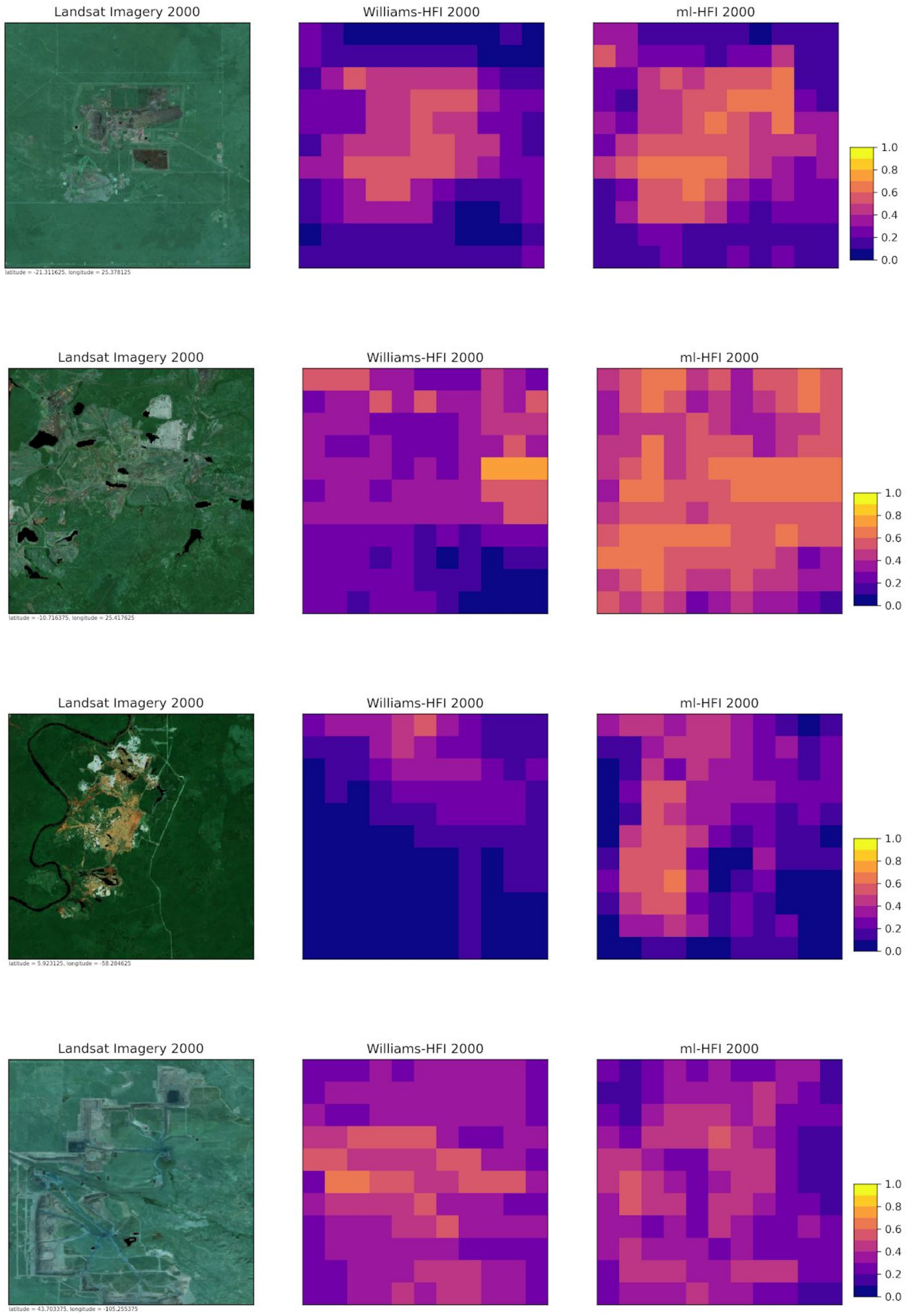


**Supp. Figure 11:** Examples of mining areas where the ml-HFI correctly predicts the human pressure on the landscape. Regions highlighted include Botswana; Democratic Republic of Congo; Guyana; USA from top to bottom.

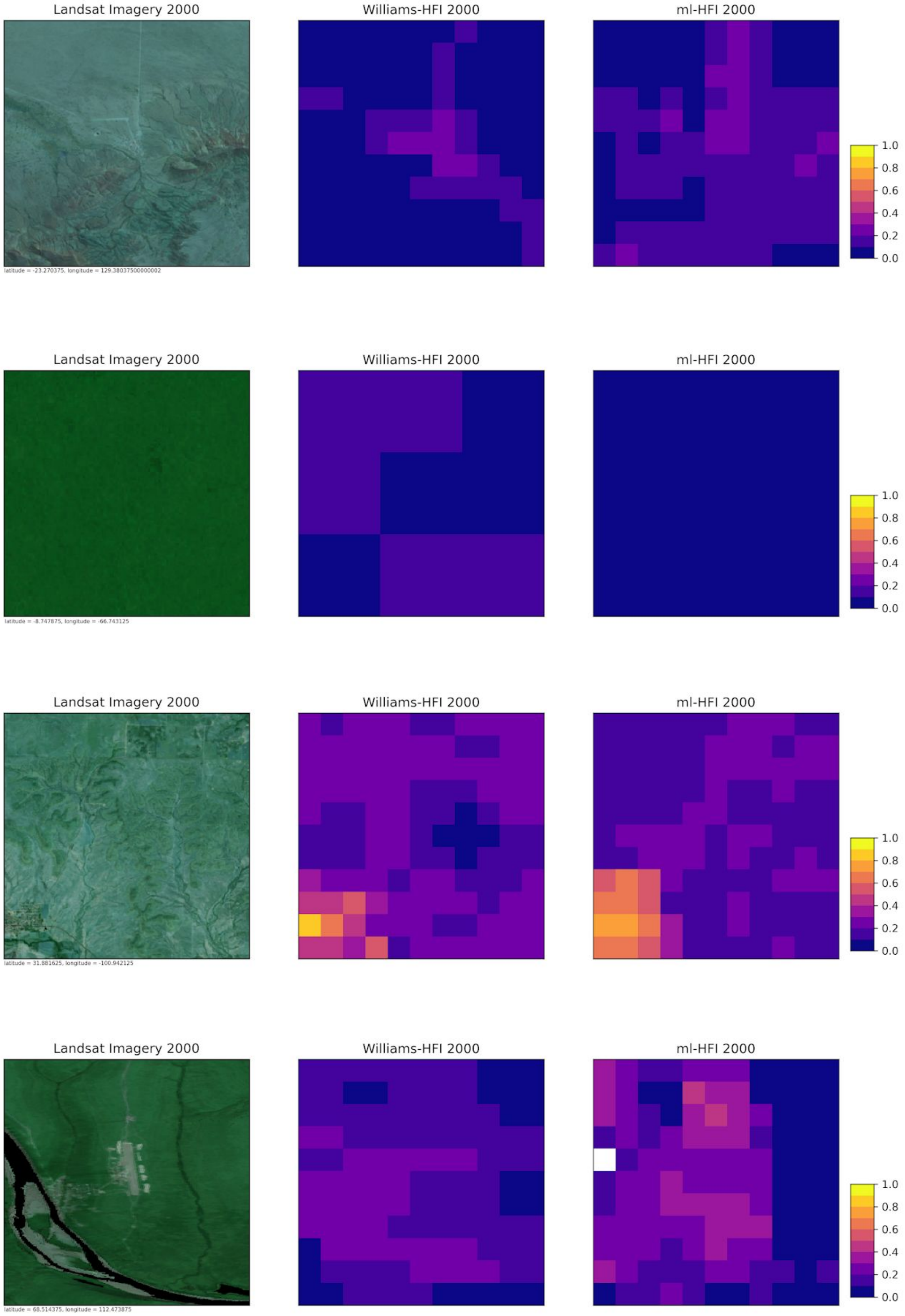


**Supp. Figure 12:** Examples of rural areas where the ml-HFI correctly predicts the human pressure on the landscape. Regions highlighted include Australia; Brazil; USA; Russia from top to bottom.

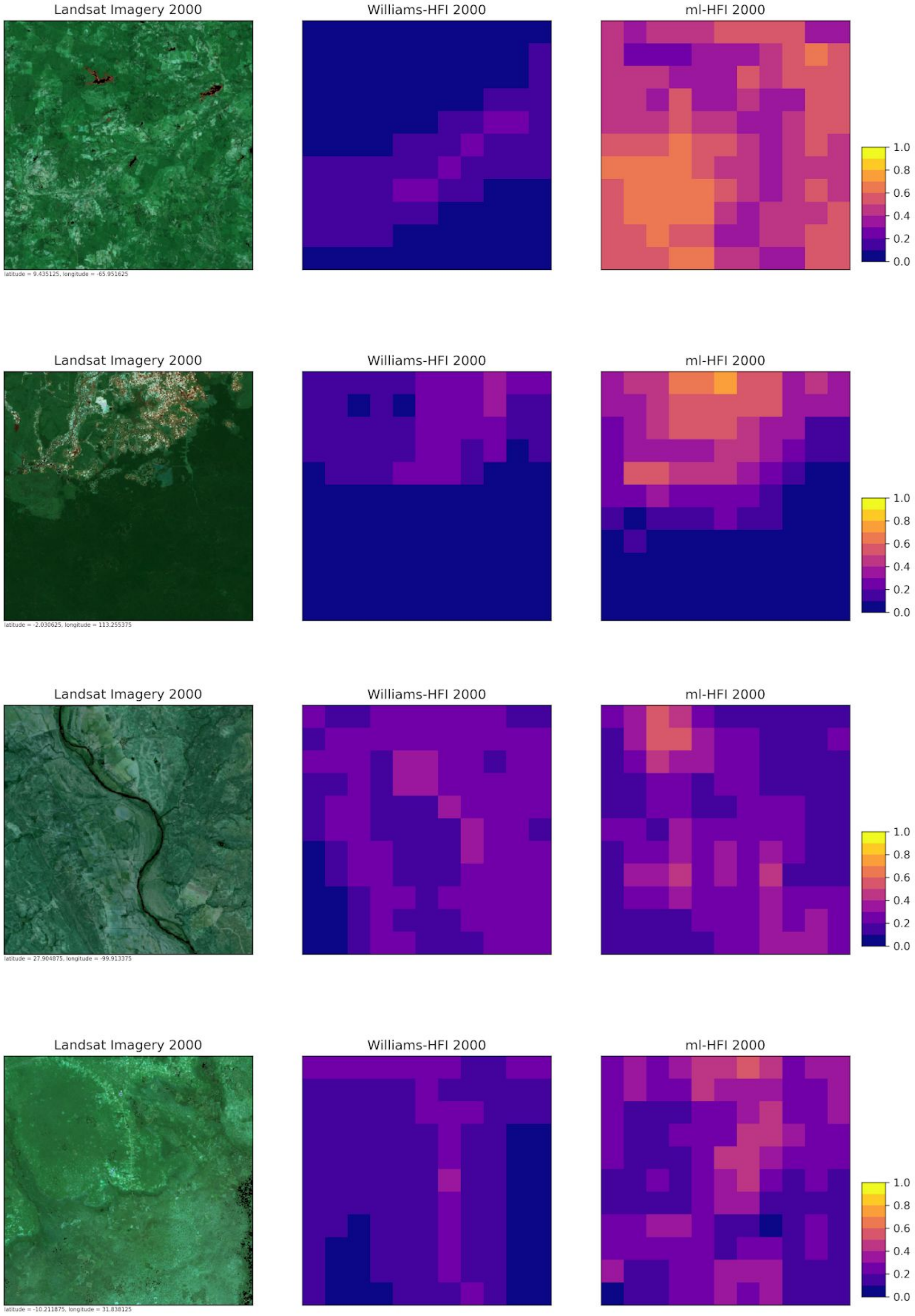


**Supp. Figure 13:** Examples of landscapes where the ml-HFI and Williams-HFI disagree and the ml-HFI appears to be correct. Regions highlighted include agriculture in Venezuela, artisanal mining in Indonesia, agriculture in Mexico, and agriculture in Zambia from top to bottom.

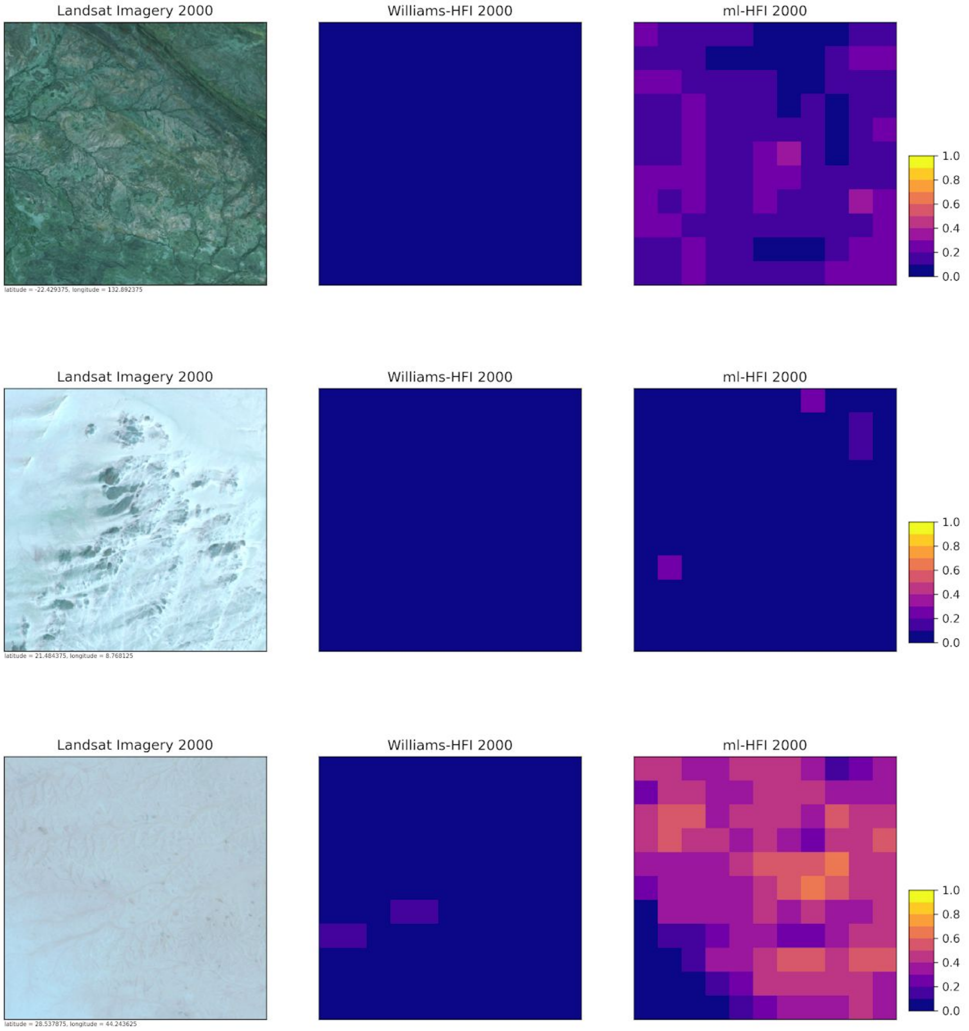


**Supp. Figure 14:** Examples of wilderness landscapes where the ml-HFI incorrectly predicts a human presence. Regions highlighted include Australia, Niger and Saudi Arabia from top to bottom.

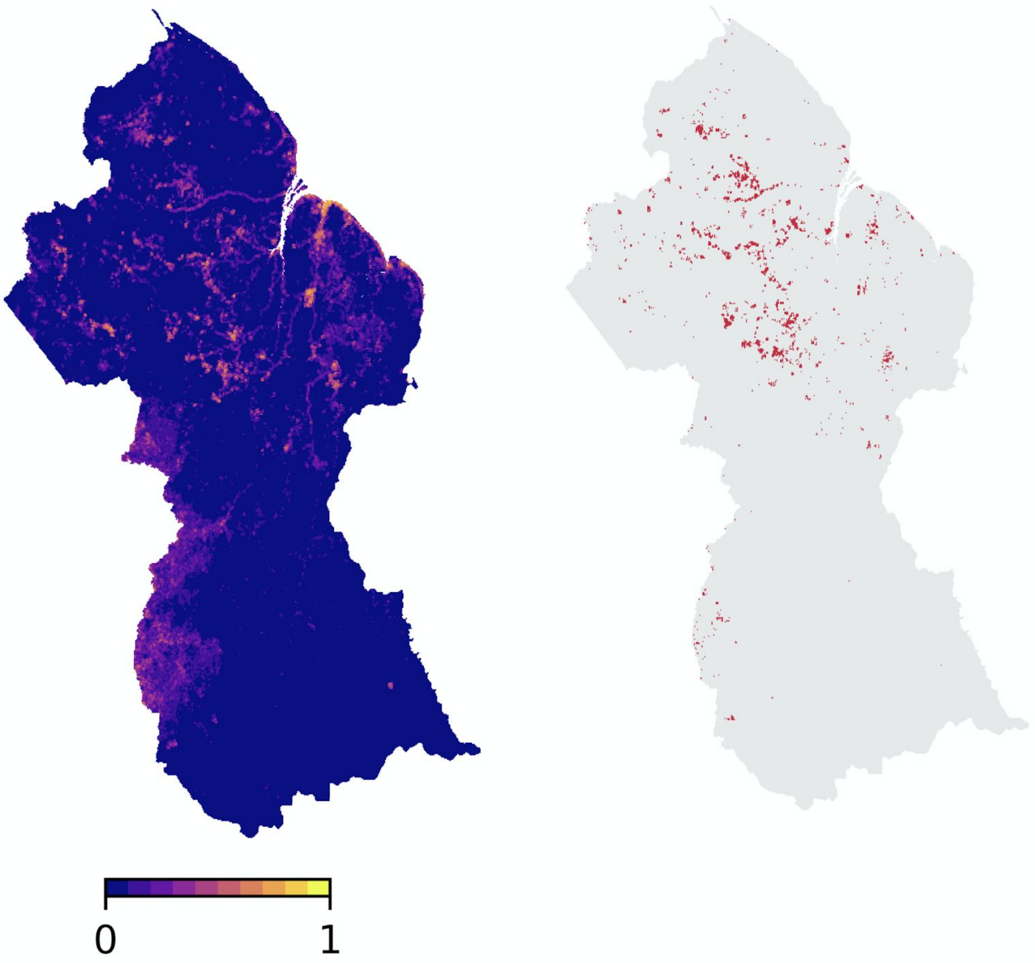


**Supp. Figure 15:** (left) ml-HFI for 2019 for Guyana. (right) Substantial increases in the ml-HFI between 2000 and 2019 in Guyana are shown in red where “substantial” refers to greater than 0.25 increase.

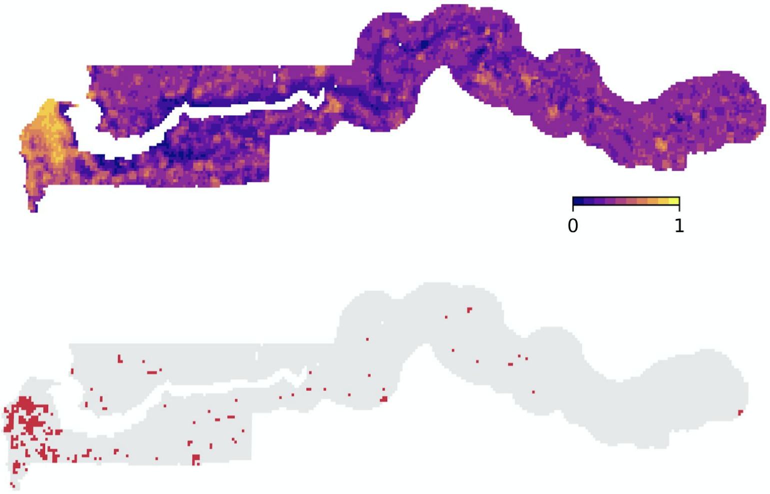


**Supp. Figure 16:** (top) ml-HFI for 2019 for The Gambia. (bottom) Substantial increases in the ml-HFI between 2000 and 2019 in The Gambia are shown in red where "substantial" refers to greater than 0.25 increase.

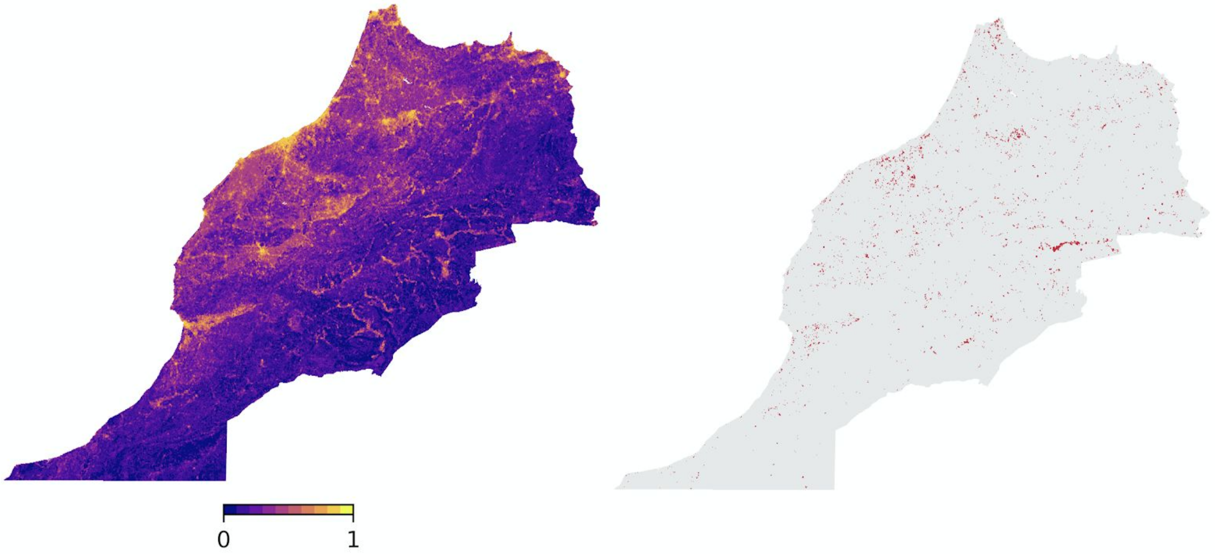


**Supp. Figure 17:** (left) ml-HFI for 2019 for Morocco. (right) Substantial increases in the ml-HFI between 2000 and 2019 in Morocco are shown in red where "substantial" refers to greater than 0.25 increase.

**Supp. Table 1:** Summary of countries that are experiencing an increase in HFI between 2000 and 2019 of greater than 0.25 across 1% or more of their land area. An SDG trend of 3 refers to ‘moderately improving’ and 4 refers to ‘on track to achieve SDG15’. The fraction of land column refers to the fraction of a country that is experiencing a change in HFI greater than 0.25. All countries in the table have been extensively quality-controlled and verified to have experienced change using the time-lapse feature on Google Earth Pro. Norway, Gabon, Libya and Sudan have been removed from the list as further inspection shows that the CNN struggles there.

| COUNTRY | SDG15 TREND | FRACTION OF LAND |
|---|---|---|
| Albania | 3 | 3% |
| Algeria | 3 | 1% |
| Andorra | 3 | 3% |
| Austria | 3 | 3% |
| Belgium | 4 | 2% |
| Benin | 4 | 2% |
| Bosnia-Herzegovina | 3 | 1% |
| Bulgaria | 4 | 1% |
| Cape Verde | 3 | 4% |
| Congo | 4 | 1% |
| Croatia | 4 | 1% |
| Cyprus | 4 | 6% |
| Czech Republic | 4 | 2% |
| Denmark | 4 | 2% |
| Dominican Republic | 3 | 2% |
| Estonia | 4 | 1% |
| Finland | 4 | 1% |
| France | 3 | 1% |
| Germany | 4 | 2% |
| Greece | 3 | 1% |
| Guinea-Bissau | 4 | 1% |
| Guyana | 4 | 2% |

| | | |
|---|---|---|
| Ireland | 4 | 3% |
| Italy | 4 | 1% |
| Liechtenstein | 4 | 10% |
| Luxembourg | 3 | 1% |
| Macedonia | 3 | 1% |
| Morocco | 3 | 2% |
| Netherlands | 4 | 5% |
| Nigeria | 3 | 1% |
| Poland | 4 | 2% |
| Portugal | 3 | 3% |
| Romania | 4 | 1% |
| Slovakia | 4 | 2% |
| Slovenia | 4 | 3% |
| Suriname | 3 | 3% |
| Sweden | 4 | 2% |
| Switzerland | 3 | 3% |
| The Gambia | 3 | 3% |
| Togo | 3 | 1% |
| Uganda | 3 | 1% |
| United Kingdom | 3 | 2% |
| Venezuela | 3 | 1% |
